# The genome sequence of the octocoral *Paramuricea clavata* – a key resource to study the impact of climate change in the Mediterranean

**DOI:** 10.1101/849158

**Authors:** Jean-Baptiste Ledoux, Fernando Cruz, Jèssica Gomez-Garrido, Regina Antoni, Julie Blanc, Daniel Gómez-Gras, Paula López-Sendino, Agostinho Antunes, Cristina Linares, Marta Gut, Tyler Alioto, Joaquim Garrabou

## Abstract

The octocoral, *Paramuricea clavata*, is a habitat-forming anthozoan with a key ecological role in rocky benthic and biodiversity-rich communities in the Mediterranean and Eastern Atlantic. Shallow populations of *P. clavata* in the North-Western Mediterranean are severely affected by warming-induced mass mortality events (MMEs). These MMEs have differentially impacted individuals and populations of *P. clavata* (i.e. varied levels of tissue necrosis and mortality rates) over thousands of kilometers of coastal areas. The eco-evolutionary processes and genetic factors contributing to these differential responses remain to be characterized. Here, we sequenced a *P. clavata* individual with short and long read technologies, producing 169.98 Gb of Illumina paired-end and 3.55 Gb of Oxford Nanopore Technologies (ONT) reads. We obtained a *de novo* hybrid assembly accounting for 712.4 Mb and 107,682 scaffolds. The contig and scaffold N50 are 15.85 Kb and 17.01 Kb, respectively. Despite of the low contiguity of the assembly, the gene completeness was relatively high, including 86% of the 978 metazoan genes contained in the metazoa_odb9 database. A total of 76,508 protein-coding genes and 85,763 transcripts have been annotated. This assembly is one of the few octocoral genomes currently available. This is undoubtedly a valuable resource for characterizing the genetic bases of the differential responses to thermal stress and for the identification of thermo-resistant individuals and populations. Overall, the genome of *P. clavata* will help to understand various aspects of its evolutionary ecology and to elaborate effective conservation plans such as active restoration actions to overcome the threats of global change.

## INTRODUCTION

The red gorgonian, *Paramuricea clavata* (Risso 1826; Figure 1), is an octocoral belonging to the Holaxonia-Alcyoniina clade (McFadden *et al*. 2006) and distributed in the Mediterranean Sea and neighboring Atlantic Ocean from 15 to 200 m depth in dim light environment (Boavida *et al*. 2016). This species plays a key ecological role as a structural species in rocky-bottoms characterized by rich diverse Mediterranean coralligenous (Ballesteros 2006). Similar to trees in terrestrial forests, *P. clavata* generates three-dimensional structures that increase overall habitat complexity which in turn has a positive impact on associated species (Ponti *et al*. 2018). This long-lived species (up to 100 years) exhibits low population dynamics: it is characterized by recruitment by pulse, a slow growth rate (mean growth rate = 0.8 cm.years^-1^), late sexual maturity (10 years of age) (Linares *et al*. 2007) and restricted dispersal and re-colonization capacities (Mokhtar-Jamaï *et al*. 2011; Arizmendi-Mejía *et al*. 2015b). The red gorgonian populations are critically impacted by defaunation, due to habitat destruction (Linares *et al*. 2007), and warming-induced mass-mortality events (MMEs) (Garrabou *et al*. 2009). Considering the biology and ecology of the species, these pressures challenge the demographic and evolutionary responses of *P. clavata*. Consequently, *P. clavata* was recently included as a vulnerable species to the IUCN red list of Anthozoans in the Mediterranean (Otero *et al*. 2017). Moreover, there is a consensus among scientists and managers regarding the urgent need to develop new resources for this species in order to promote its conservation.

**Figure 1.**
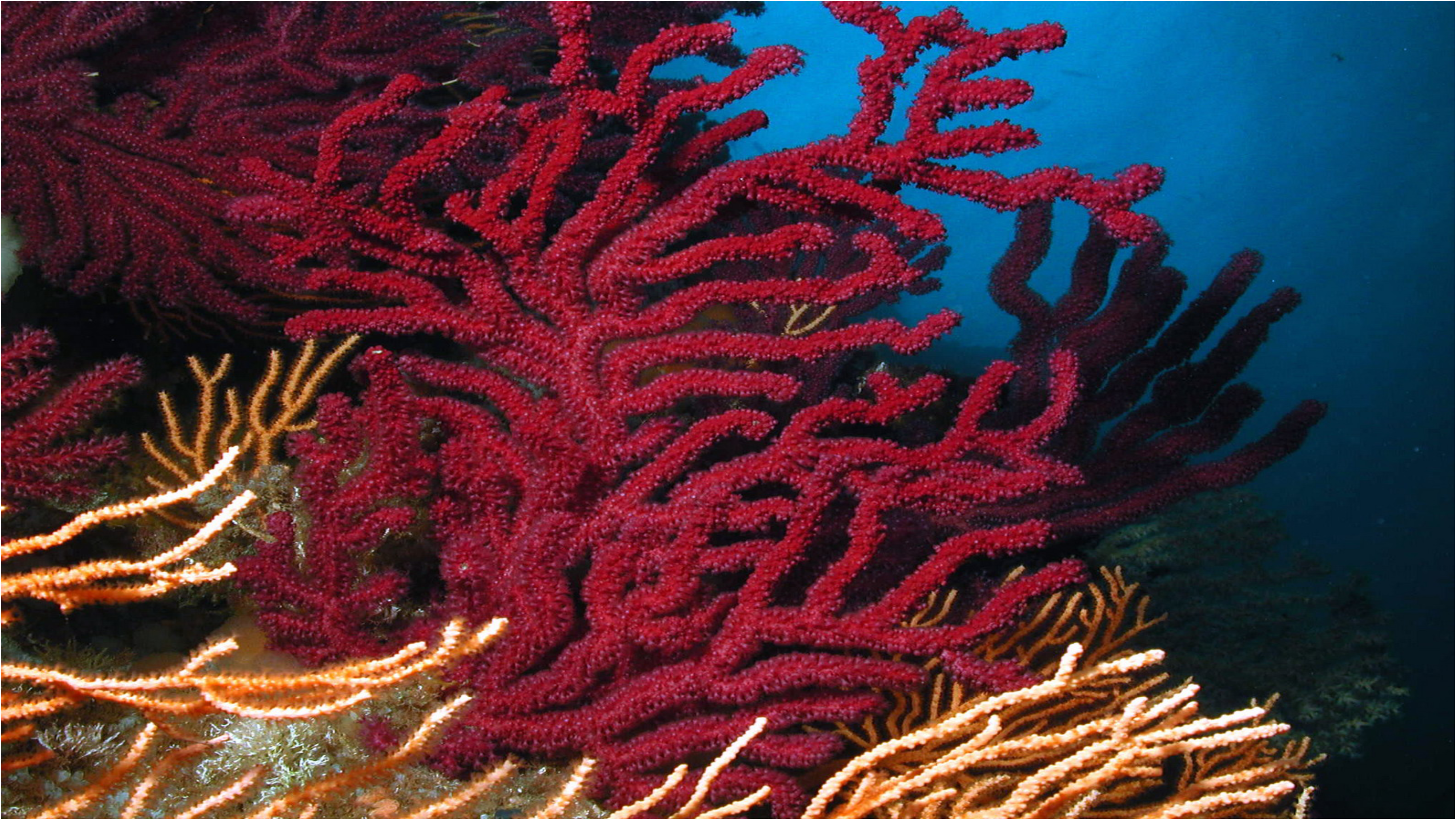
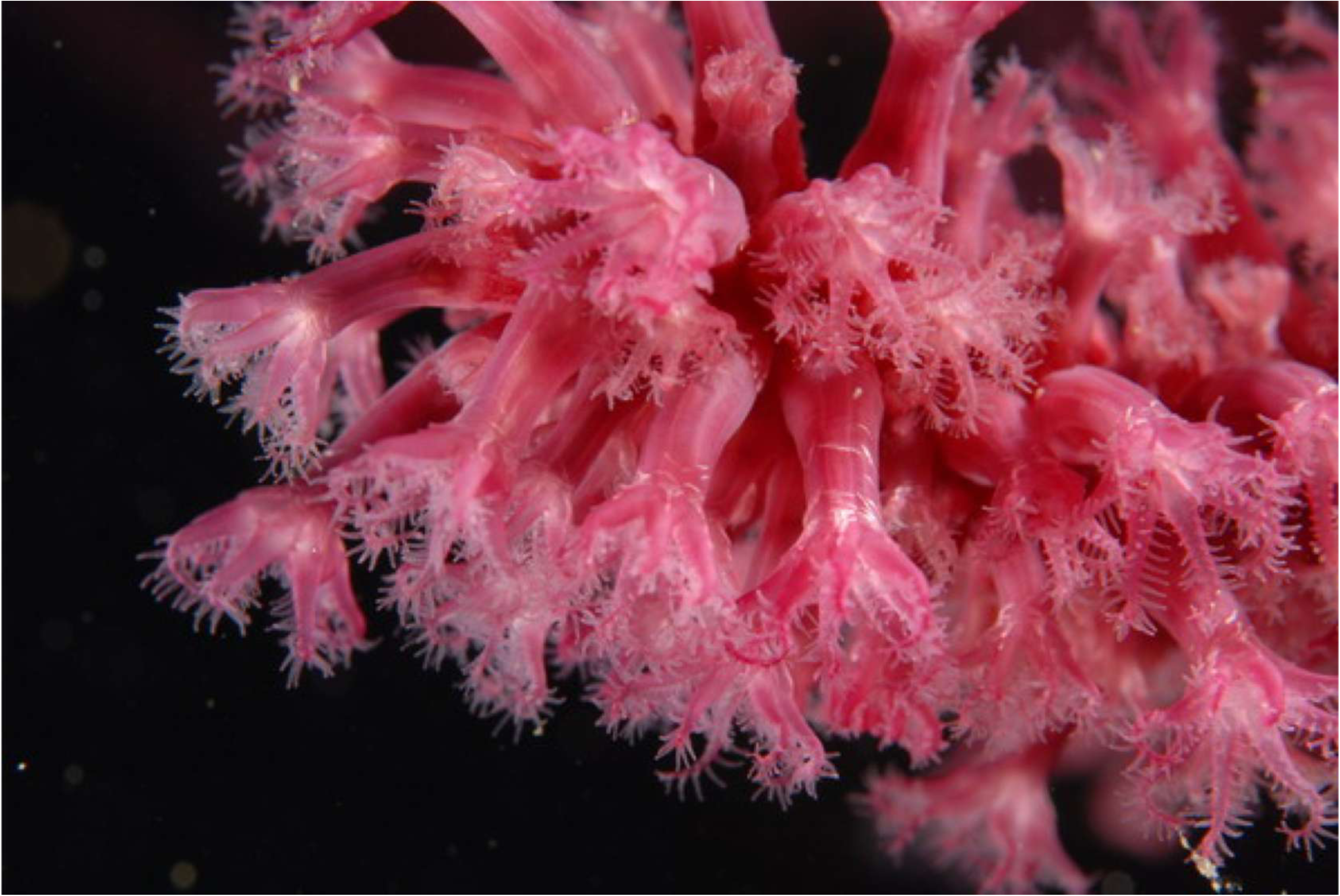
The red gorgonian *Paramuricea clavata* (Risso, 1826): a) whole colony; b) close up on the polyps.

Focusing on mass-mortality events (MMEs), intensive field surveys have demonstrated the differential impact of warming on individuals and populations of *P. clavata*. For instance, during the 2003 MME, the percentage of affected colonies (i.e. showing tissue necrosis) ranged from less than 5% up to more than 80% depending on the population (Garrabou *et al*. 2009). Thus, individuals and/or populations show different levels of tolerance to thermal stress, suggesting the occurrence of warming-resistant individuals. The presence of these individuals affords a new perspective for the conservation of the species, especially in terms of active restoration actions.

Accordingly, population-by-environment interactions (PEI) focused on the interactions with thermal environment have been receiving more attention in *Paramuricea clavata*. Common garden experiments in controlled conditions have been used to identify different physiological factors (e.g., sex, sexual maturity) driving the differential responses to thermal stress reported from the field (Coma *et al*. 2009; Arizmendi-Mejía *et al*. 2015b). In the meantime, different studies have aimed to decipher the respective role of selected (local adaptation) and neutral (genetic drift) eco-evolutionary processes on the resistance to thermal stress (Ledoux *et al*. 2015; Crisci *et al*. 2017). While genetic drift seems to play a central role in *P. clavata* PEI, definitive conclusions regarding the eco-evolution of thermo-resistance are still lacking mainly because of the limited genetic tools used (e.g., low number of genetic markers). In order to promote the conservation of *Paramuricea clavata*, we aim to develop genomic resources to gain insights into the eco-evolutionary processes and genetic factors involved in the differential response to thermal stress.

## METHODS & MATERIALS

### Sample Collection

One apical tip (8 cm) of a reproductive colony (> 30 cm) of *P. clavata* was sampled by SCUBA diving at 20m depth in the Cova de la Vaca (42°2’52.97’’N; 3°13’34.76’’E) in The Montgrí, Medes Islands and Baix Ter Natural Park (Catalunya, Spain). The sample was transferred alive to the Experimental Aquaria Facility (ZAE) of the Institute of Marine Science (Barcelona; Spain) and placed in a 70L tank filled with filtered Mediterranean Sea water pumped from 10m depth in a continuous flux system. This sample was divided in three fragment sections (3_10-10, 5_27-10 and 6_2-1) for the DNA extractions described below.

### Genomic DNA extraction

Total genomic DNA was extracted from fresh tissue frozen in liquid nitrogen using the Gentra PureGene Tissue Kit (Qiagen) following manufacturer protocol. DNA purity and quantity were estimated using spectrophotometer and Qubit fluorescent based kit (Thermo Fisher Scientific). DNA integrity was assessed on 0.8% agarose gel electrophoresis.

### Whole Genome Sequencing with Illumina

The Roche-Kapa Biociences kit for short-insert paired-end libraries for Illumina was used for DNA library preparation of *P. clavata* with some minor modifications. A pool of seven gDNA extractions from fragment section 3_10-10 was re-purified with AMPure XP Beads (Agencourt, Beckman Coulter) and eluted in 50ul of water. Genomic DNA (6.0 μg) was sheared on a Covaris™ LE220 in order to reach DNA fragment sizes of ∼400-800bp. The fragmented DNA was size-selected on 1% agarose gel where eight bands were excised to isolate DNA fragments of precise insert size (520bp). Three gel fractions were selected for further purification with Qiagen QIAquick® Gel Extraction Kit and the size was determined on an Agilent 2100 Bioanalyzer with the DNA7500 assay (362bp, 429bp, 548bp, fractions D, E, F), end-repaired, adenylated and ligated to dual matched indexed paired-end adaptors (IDT). The adaptor ligated library size (458bp, 516bp, 678bp) was confirmed on an Agilent 2100 Bioanalyzer with the DNA7500 assay. All libraries were quantified with the Library Quantification Kit for Illumina Platforms (Roche-Kapa Biosystems). The sequencing library with the mean insert size of 429bp (agarose gel, fraction E) was sequenced using TruSeq Rapid SBS Kit v2 (Illumina), in paired end mode, 2×251bp, in two sequencing lanes of Illumina HiSeq2500 flowcell v2 (Illumina) according to standard Illumina operating procedures with a minimal yield of 170 Gb of raw data. Primary data analysis, image analysis, base calling and quality scoring of the run, were processed using the manufacturer’s software Real Time Analysis (RTA 1.18.66.3) and followed by generation of FASTQ sequence files by CASAVA.

### Long Read Whole Genome Sequencing

Genomic DNA of *P. clavata* was obtained from four extractions of fragment section 5_27-10 and six extractions from fragment section 6_2-1. All extractions from each respective fragment section were then pooled into two samples (pooled sample 5_27-10 and pooled sample 6_2-1) and re-purified using AMPure XP Beads (Agencourt, Beckman Coulter) adding 0.4 volume (V/V) to the pooled sample. Both pooled samples were quality controlled using Pippin Pulse (Sage Science) and with Nanodrop (Thermo Fisher Scientific) ratios 260/230 and 260/280. From each pooled sample a sequencing library was constructed using the Ligation Sequencing Kit 1D, SQK-LSK108 (Oxford Nanopore Technologies) starting with 2μg of restricted integrity gDNA without a fragmentation step. The DNA was repaired using the NEBNext FFPE Repair Mix (New England Biolabs), end-repaired and adenylated with the NEBNext Ultra II End Repair and A-Tailing Module (New England Biolabs) and MinION AMX adapters (Oxford Nanopore Technologies) were ligated using the NEB Blunt/TA Ligase Master Mix (New England Biolabs). Each step was followed by purification with AMPure XP Beads. The DNA/beads ratio was 1 (V/V) after the end-repair and adenylation step. After the repair and the final purification/size selection steps the DNA/beads ratio was 0.4 (V/V) in order to eliminate all fragments below 2kb.

The sequencing run was performed on a MinION instrument (Oxford Nanopore Technologies) using the R9.5 chemistry FLO-MIN107 flowcell for the first run (pooled sample 5_27-10) and the R9.4.1 chemistry FLO-MIN106 flowcell (Oxford Nanopore Technologies) for the second run (pooled sample 6_2-1), according to manufacturer’s recommendations. In brief, first the MinKNOW interface QC (Oxford Nanopore Technologies) was run in order to assess the flowcell quality and followed by flowcell priming. The sequencing library was mixed with running buffer, Library Loading Beads (Oxford Nanopore Technologies) and nuclease free water and loaded onto the “spot on” port for sequencing. The sequencing data was collected for 48 hours. The quality parameters of the sequencing runs were further monitored by the MinKNOW 1.10.16 platform while the run was base-called using the Albacore v2.0.1 agent in real time.

### Genome size and complexity

As there is no empirical estimate for *P. clavata* genome size, we downloaded 41 C-value estimates corresponding to the Cnidaria phylum in the Animal Genome Size Database (Gregory 2017). In addition, we ran two different k-mer analyses on the raw PE reads to estimate the size and complexity of the genome. First, we examined the frequency distribution of k-mers, 57 bp long, using Jellyfish v2.2.6 (Marçais and Kingsford 2011) and then ran GenomeScope v.1.0 (Vurture *et al*. 2017). An additional estimate, was obtained with SGA preqc (Simpson 2014) (using a k=31).

### Filtering contaminated reads and trimming PE reads to 150 bp

Before de novo assembly, all reads from PE400 were filtered out from contaminants by mapping (gem-mapper (Marco-Sola *et al*. 2012) with up to 2% mismatches) against a contamination database that included phiX, Univec sequences, *E. coli* and various contaminants detected with kraken (Wood and Salzberg 2014) in more than 0.01% of the reads (see Table 3). Note that our read decontamination method is stringent enough to remove real contaminants (almost exact matches with ≤ 2% mismatches) such as phiX but does not detect similar but divergent sequences (such as different bacterial strains); these are detected in the final assembly by using BLAST or during the genome annotation.

The filtered Illumina PE were trimmed to 150bp using FASTX toolkit v.0.0.13 (http://hannonlab.cshl.edu/fastx_toolkit/). This was done to optimize the de *Bruijn* graph construction. The trimming reduced the sequencing coverage to 142.12x (close to 120x – the ideal for a heterozygous genome) and increased the mean base quality of the reads (last cycles produce lower base qualities).

### *De novo* hybrid assembly

Two different hybrid assemblies were obtained with MaSuRCA v3.2.6 [21, 22]: one using the complete PE400 2×251bp library (pcla1) and a second using the reads trimmed to 150bp (pcla4). In both cases, the reads were assembled with CELERA and the USE_LINKING_MATES option. As part of our genome annotation strategy (see below), we also removed 2,332 scaffolds contaminated with bacteria or fungi from the most contiguous hybrid assembly (see below). In addition, we mapped the complete mitochondrial genome (NC_034749.1) to the decontaminated assembly using minimap2 v2.14 (Li 2018).

### RNA Library Preparation and Sequencing

To obtain a transcriptome for annotation, total RNA was extracted from three different individuals coming from three distant populations (LaVaca, Gargallu and Balun) from three regions (Catalan Sea, Corsica and Croatia) and submitted to a thermal stress in controlled conditions in aquaria. Sampling was conducted by scuba-diving around 20m depth in each locality. For each individual, an 8 cm apical tip was sampled (hereafter colony) and placed in coolers with seawater ice packs to maintain the water temperature between 15–18 °C. The colonies were transported alive to experimental aquaria facilities at the Institute of Marine Sciences-CSIC (Barcelona, Spain). The maximum transportation time was of 36 h for the colonies collected in Gargallu (Corsica, France) and Balun (Eastern Adriatic, Croatia). Until the beginning of the experiment, colonies were maintained at control temperatures (16–17 °C) presenting expanded polyps during feeding events and no tissue necrosis indicating their healthy conditions. The experiment set-up was inspired from previous experiments with the same species [13, 27]. Briefly, after an acclimation period, the seawater was heated to 25°C during 24h and maintained for 25 days. The tank was equipped with submersible pumps to facilitate water circulation. Temperature was registered with Tidbit Stowaway autonomous temperature sensors every half an hour. The experimental set functioned as an open system. Tissues were sampled and conserved in RNA later ^TM^ (Qiagen) for each individual before heating (T0) and after 25 days of thermal stress (T1). RNA extractions were conducted combining TRI Reagent Solution (Invitrogen) for tissue lysis and phase separation and Qiagen RNeasy Mini protocol for purification and elution. RNA extractions were pooled in two groups: one including the extractions at T0 and one including the extractions at T1 and quantified by Qubit® RNA BR Assay kit (Thermo Fisher Scientific). RNA integrity was estimated by using the RNA 6000 Nano Bioanalyzer 2100 Assay (Agilent).

The RNASeq libraries were prepared from total RNA using KAPA Stranded mRNA-Seq Kit Illumina® Platforms (Roche-Kapa Biosystems) with minor modifications. Briefly, after poly-A based mRNA enrichment with oligo-dT magnetic beads and 500ng of total RNA as the input material, the mRNA was fragmented. The strand specificity was achieved during the second strand synthesis performed in the presence of dUTP instead of dTTP. The blunt-ended double stranded cDNA was 3’adenylated and Illumina platform compatible adaptors with unique dual indexes and unique molecular identifiers (Integrated DNA Technologies) were ligated. The ligation product was enriched with 15 PCR cycles and the final library was validated on an Agilent 2100 Bioanalyzer with the DNA 7500 assay.

The libraries were sequenced on HiSeq 4000 (Illumina, Inc) with a read length of 2×76bp using HiSeq 4000 SBS kit in a fraction of a HiSeq 4000 PE Cluster kit sequencing flow cell lane generating a mean of 80 million paired end reads per sample. Image analysis, base calling and quality scoring of the run were processed using the manufacturer’s software Real Time Analysis (RTA 2.7.7).

### Genome annotation

First, repeats present in the pcla4 genome assembly were annotated with RepeatMasker v4-0-6 (http://www.repeatmasker.org) using the repeat library specific for our assembly that was built with RepeatModeler v1.0.11. Repeats that were part of repetitive protein families (detected by running a Blast of the Repeat library against swissprot) were removed from the library before masking the genome.

An annotation of protein-coding genes was obtained by combining RNA-seq alignments, protein alignments and *ab initio *gene predictions. A flowchart of the annotation process is shown in Figure 6.

RNA-seq reads were aligned to the genome with STAR v-2.6.1b (Dobin *et al*. 2013) and transcript models were subsequently generated using Stringtie v1.0.4 (Pertea *et al*. 2015). PASA v2.3.3 (Haas *et al*. 2008) was used to combine the Stringtie transcript models with 534 soft coral nucleotide sequences downloaded from NCBI in January 2019. The *TransDecoder* program, which is part of the PASA package, was run on the PASA assemblies to detect coding regions in the transcripts. Then, the complete *Stylophora pistilata* proteome was downloaded from Uniprot (January 2019) and aligned to the genome using SPALN v2.3.1 (Gotoh 2008). *Ab initio* gene predictions were performed on the repeat-masked pcla4 assembly with four different programs: GeneID v1.4 (Parra *et al*. 2000), Augustus v3.2.3 (Stanke *et al*. 2006), GlimmerHMM (Majoros *et al*. 2004) and Genemark-ES v2.3e (Lomsadze *et al*. 2014) with and without incorporating evidence from the RNA-seq data. All the gene predictors except Genemark, which runs in a self-trained manner, were run with the parameters obtained by training with a set of high-quality candidate genes extracted from the Transdecoder results. Finally, all the data was combined into consensus CDS models using EvidenceModeler-1.1.1 (Haas *et al*. 2008)). Additionally, UTRs and alternative splicing forms were annotated through two rounds of PASA annotation updates. The resulting transcripts were clustered into genes using shared splice sites or significant sequence overlap as criteria for designation as the same gene. Functional annotation of the annotated proteins was done using Blast2go (Conesa *et al*. 2005), which in turn ran a BLASTP (Altschul *et al*. 1997) search against the non-redundant database (March 2019) and Interproscan (Jones *et al*. 2014) to detect protein domains on the annotated proteins.

Finally, the annotation of non-coding RNAs (ncRNAs) was performed as follows. First, the program cmsearch (v1.1) (Cui *et al*. 2016) that comes with Infernal (Nawrocki and Eddy 2013) was run against the RFAM (Nawrocki *et al*. 2015) database of RNA families (v12.0). Also, tRNAscan-SE (v1.23) (Lowe and Eddy 1997) was run in order to detect the transfer RNA genes present in the genome assembly. PASA-assemblies longer than 200bp that had not been annotated as protein-coding and not overlapped by more than 80% by a small ncRNA were incorporated into the ncRNA annotation as long-non-coding RNAs (lncRNAs).

Thanks to the functional annotation, we were able to detect the presence of bacterial and fungal genes, suggesting some residual contamination in the assembly (pcla 4). Therefore, we removed potential contaminant sequence by combining as criteria for retention the gene functional annotation, the mean GC content and the presence of expression and *P. clavata*-specific repeats for each scaffold. As a result of this decontamination process, 2,322 scaffolds were removed from the assembly as they belonged mainly to *Aspergillus*, *Endozoicomonas* or other bacteria. The gene completeness of the pcla6 assembly (i.e. decontaminated and free of mitochondrion) was estimated with BUSCO v3 (Simão *et al*. 2015) using the metazoa database of 978 conserved genes.

### Genome-wide Heterozygosity (SNVs)

Re-sequencing at enough depth (>20x) allows extracting valuable genome-wide information from a single diploid sample by simply re-mapping against the reference genome and calling variant. Although adaptor removal and quality trimming are not recommended for MaSuRCA, they are strictly necessary before variant calling. Therefore, we detected and trimmed Illumina adaptor sequences and performed quality trimming in PE400. For this purpose, we used the Trim Galore! wrapper script (http://www.bioinformatics.babraham.ac.uk/projects/trim_galore/) with -q 10 and then we filtered out contaminated reads, as described above (Table 3). Finally, all these reads were mapped against pcla6 using BWA MEM v.0.7.7 (Wood and Salzberg 2014) and the –M option to discard mappings of chimeric reads.

For variant calling, we used GATK 3.7 (McKenna *et al*. 2010; DePristo *et al*. 2011), adapting the “GATK Best Practices HaplotypeCaller GVCF” (Van der Auwera *et al*. 2013) to a diploid organism without a set of known variants such as the Single Nucleotide Polymorphism database (dbSNP). Specifically, we did not perform base quality score recalibration, as the model normally degrades the base qualities in the absence of known variant sites. First, the BWA alignment was screened for duplicates using MarkDuplicates of PICARD v1.6 (https://broadinstitute.github.io/picard/). Then, we identified the callable sites per sample using the GATK’s CallableLoci tool, with options --minBaseQuality 10 -- minMappingQuality 20, to be in concordance with the default value for these parameters in HaplotypeCaller (see below). The actual variant calling was performed using the HaplotypeCaller but restricting it to callable sites and with options: -dt NONE -rf BadCigar -- never_trim_vcf_format_field -ploidy 2 --min_base_quality_score 10 -- standard_min_confidence_threshold_for_calling 30 --emitRefConfidence GVCF and -- GVCFGQBands at Genotype Qualities 15, 20, 25, 30, 35, 40, 45, 50, 55, 60, 65, 70, 80 and 99. The resulting GVCF was used to call genotypes with the GenotypeGVCF tool and option --never_trim_vcf_format_field.

After variant calling, we exclusively considered supported SNVs, defined as those that are bi-allelic, covered at least by 10 reads and at least two reads supporting the alternative allele at heterozygous sites. The kmer analyses (see results; Figure 4) shows that we have artificially duplicated some sequences (k-mers), due to an inefficient collapse of alternative haplotypes, in both the homozygous peak and the heterozygous peak. These duplicated sites may contribute to an underestimation of heterozygosity because the reads corresponding to each allele can potentially map separately to two independent locations in the assembly rather than to the same locus. For this reason, we extracted all 57-mers repeated exactly twice in the heterozygous and homozygous peaks of the assembly (pcla6). Our approach used first JELLYFISH v2.2.6 to dump all 57-mers contained in the PE reads (with coverage between 20x and 200x, see Figure 4) into a FASTA file. Second, KAT v2.3.3 (Mapleson *et al*. 2017) (kat sect with options -F -G 2) to get all 57-mers contained in the FASTA repeated exactly twice. The third step consisted in obtaining the location of these artificially duplicated 57-mers by looking for exact mappings with GEM-mapper build 1.81 (parameters -e 0 -m 0 -s 0). Afterwards, these genomic intervals were merged into a BED file (using BEDTools v.2.16.2 (Quinlan and Hall 2010)) and subtracted from the callable sites with BEDOPS/2.0.0a (Neph *et al*. 2012). Finally, we selected all SNV variants at callable sites that do not contain artificially duplicated 57-mers using GATK v.3.7.

### Microsatellite Markers

Previous molecular markers were identified and used for the species without information about their exact location in the genome. The newly sequenced and annotated genome has provided the means to do this. In order to know the genomic context around these markers, we mapped the 18 available microsatellite markers (Mokhtar-Jamaï *et al*. 2011; Ledoux unpublished) to pcla6 using our BLAST server (http://denovo.cnag.cat/genomes/pclavata/blast/).

## RESULTS AND DISCUSSION

### Whole Genome Sequencing with Illumina and Nanopore sequencing technologies

The Illumina HiSeq2500 run produced 338.6 million pairs of 251bp reads, accounting for a total of 169.98 Gb of sequence, representing 239x coverage of the genome (Table 1). The total nanopore yield accounted for 3.55 Gb (∼5x coverage) and read lengths were relatively low, with a read N50 of 2.67 Kb instead of the expected 7-12Kb. It is noteworthy that the efficiency of the nanopore sequencing was heavily influenced by the quality of the DNA extraction.

**Table 1.**
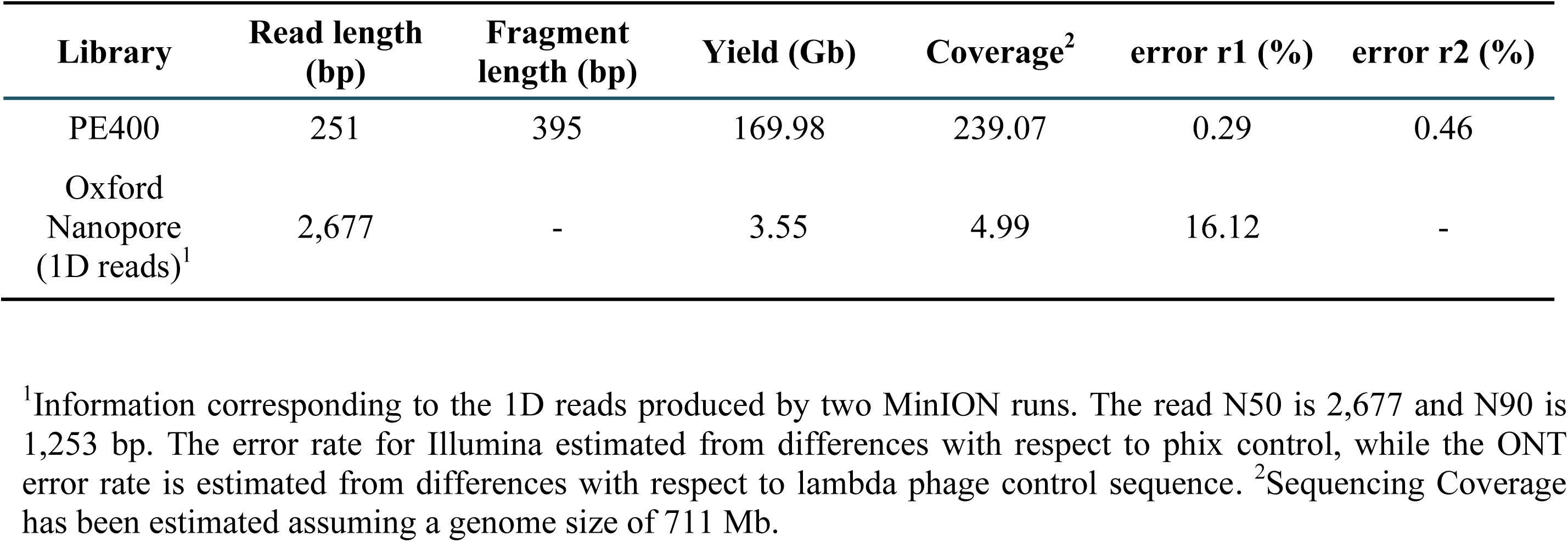
Whole Genome Sequencing Output.

### Genome size and complexity

We show the results of two different k-mer analyses and compare them with the empirical data available for the phylum Cnidaria. GenomeScope fits the 57-mer profile to a mixture model (Model Fit: 92.52-96.72%). This method estimates the haploid genome size to be between 711.53 and 712.31 Mb (Table 2). The analysis also suggests that the genome possess an appreciable amount of heterozygosity (0.91%) and high repetitiveness (41.5% of the genome is likely to be repeated) (Figure 2 and 3). The GenomeScope estimate is close to the estimate of 747.45 Mb obtained with SGA preqc (using a k=31).

**Figure 2.**
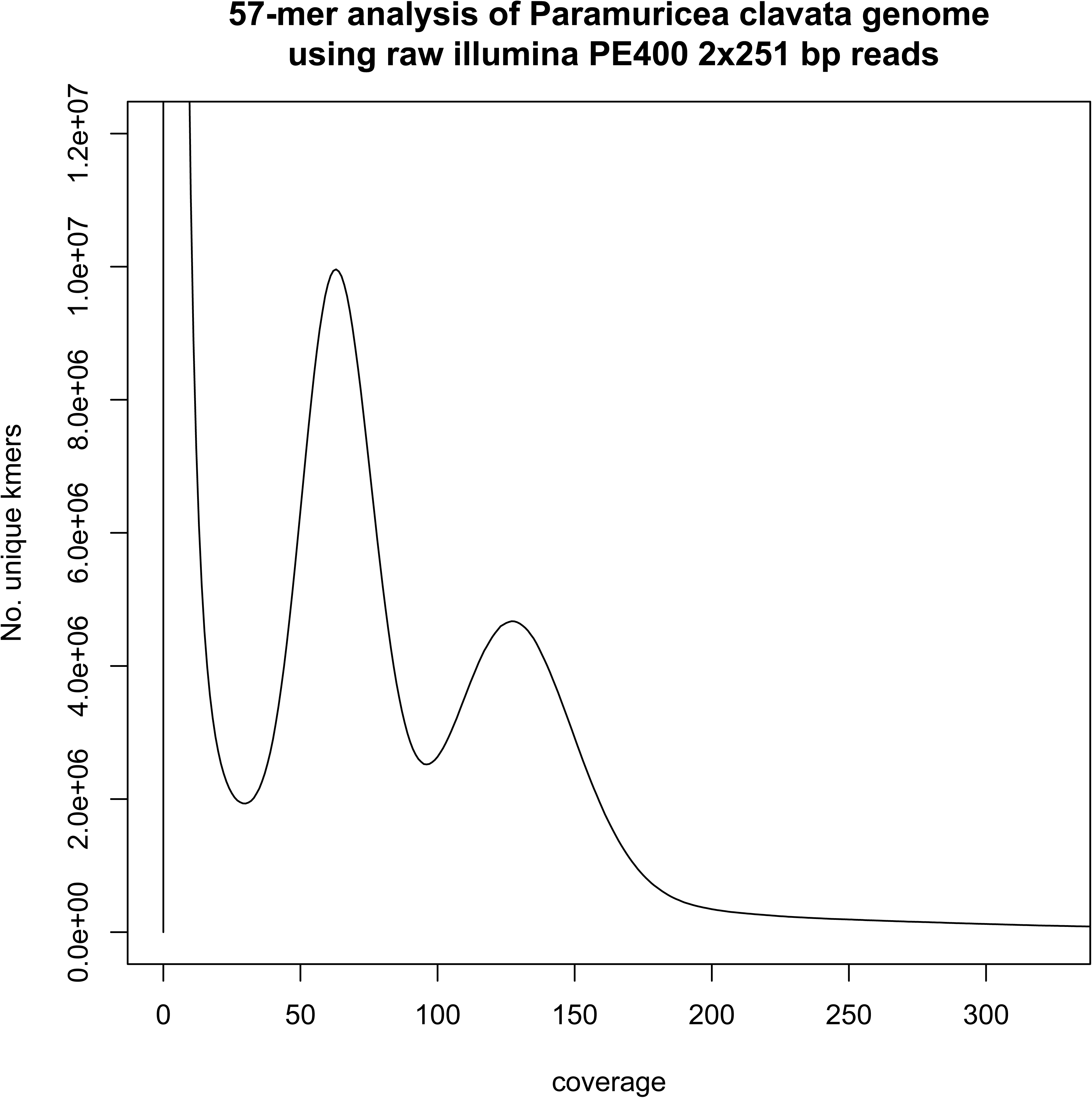
57-mer analysis of the sequenced genome. All 57-mers in the PE400 library were counted and the number of distinct 57-mers (kmer species) for each depth from 1 to 250 are shown in this plot. The main homozygous peak at depth 124 corresponds to unique homozygous sequence and a tall heterozygous peak lies at half depth of it (62). Finally, the thick long tail starting at depth 180 corresponds with repetitive k-mers in the genome. The high peak at very low depths, caused by sequencing errors, has been truncated to facilitate representation.

**Figure 3.**
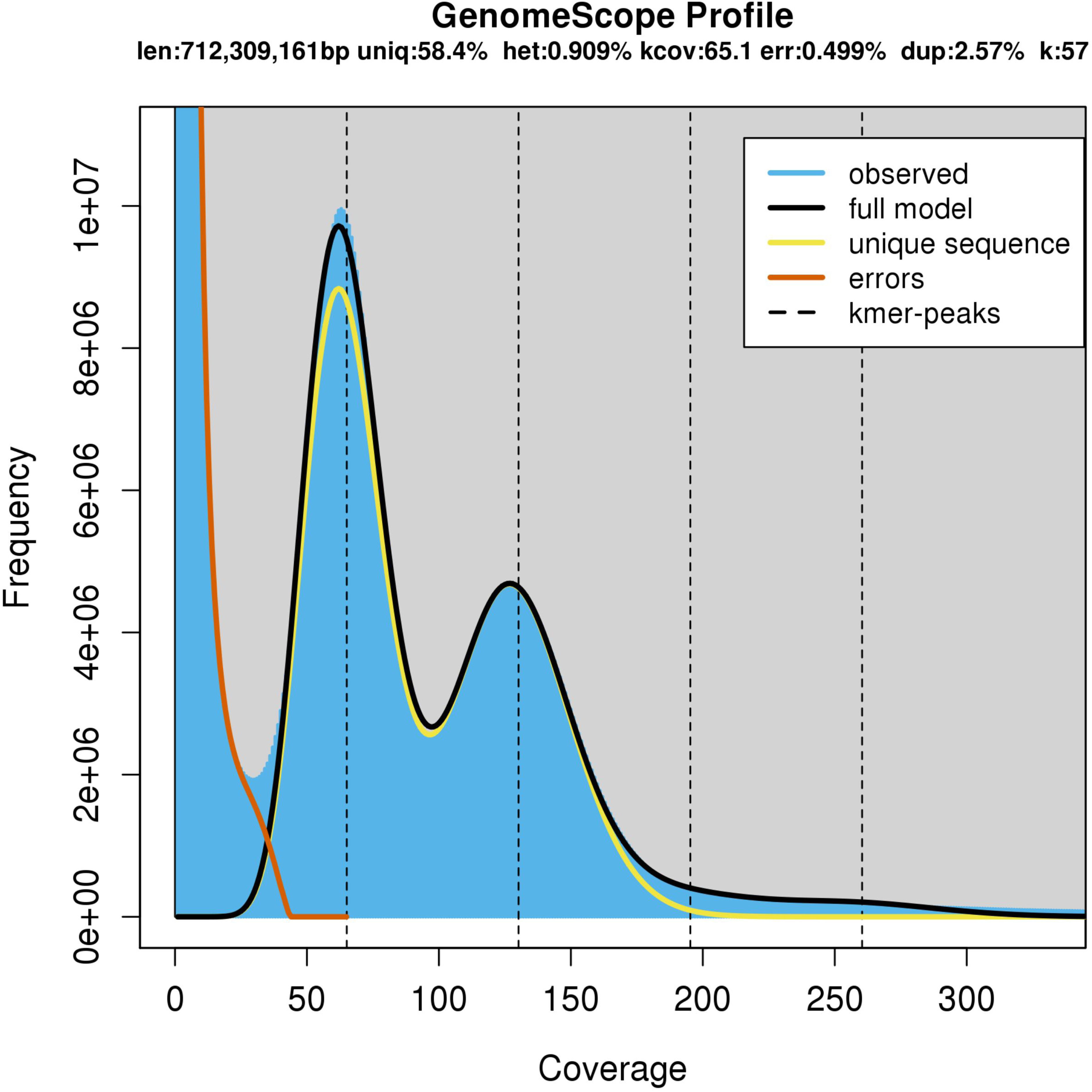
k-mer profile and model fit as estimated with GenomeScope v.1.0 from PE400 using a k-mer length of 57 bp. Note that the model finds an excess of repetitive sequence in the rightmost tail of the distribution after approximately 180 bp.

**Table 2.**
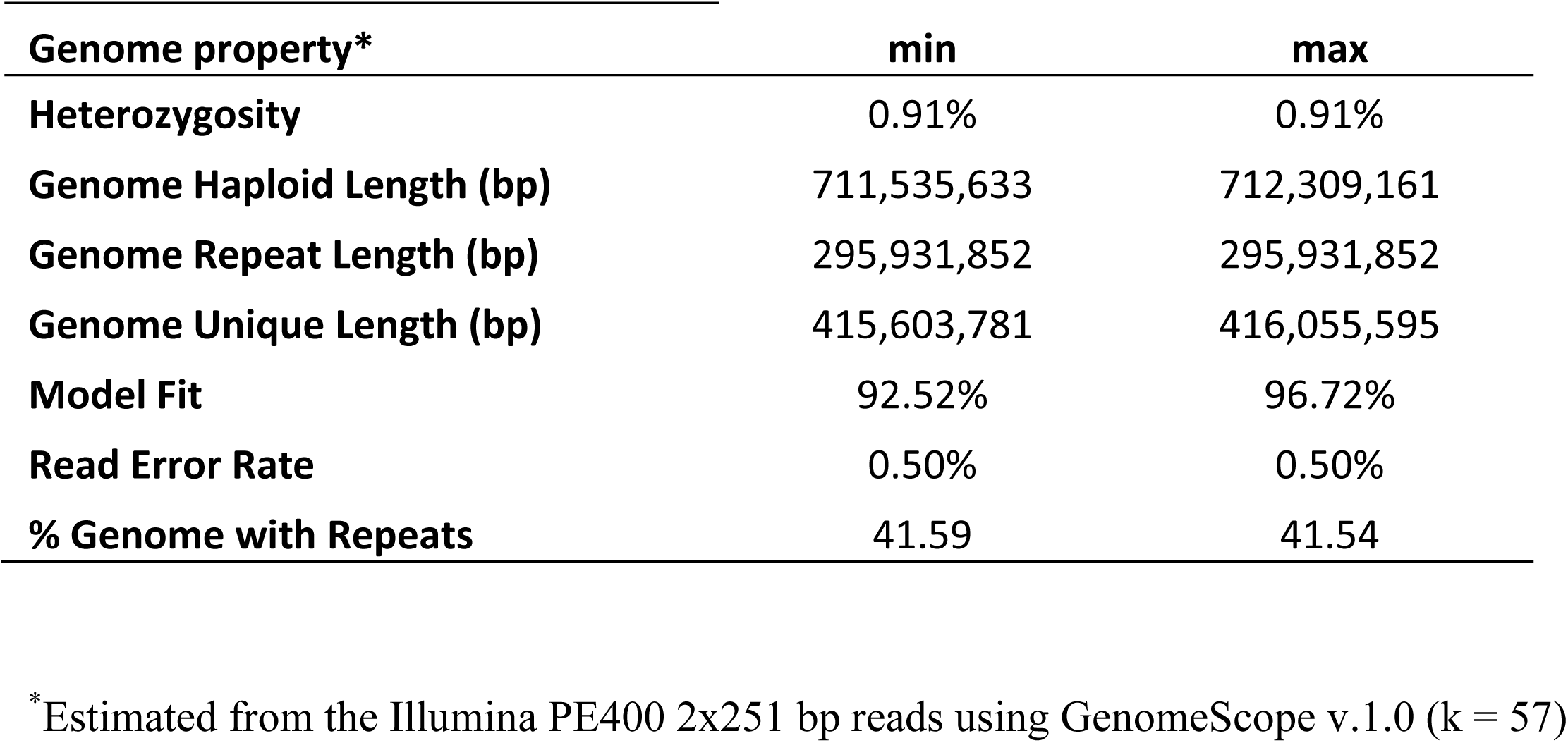
Genome Properties.

The genome size of cnidarians shows large variations, ranging from just 224.92 Mb in *Nematostella vectensis* (T.R. Gregory, unpublished data) to the 3.48 Gb estimated for *Agalma elegans* (Adachi *et al*. 2017). However, the mean cnidarian genome size is 838.45 Mb and the closest estimate to *P. clavata* belongs to *Sarcophyton sp*. (another Antozoan member of the Order Alcyonacea), that has a haploid genome size of 625.94 Mb (Adachi *et al*. 2017) supporting the value of 711.53 Mb as a reliable estimate of the genome size of *P. clavata*.

### *De novo* hybrid assembly

The more contiguous assembly was obtained using 142x coverage with the 2×150 bp trimmed reads and 5x of ONT long-read data (i.e. pcla4). We interpret this in two different but related ways. First, trimming 100bp off of the reads results in a reduction of the number of error k-mers that complicate the construction of the de *Bruijn* graph. Second, de *Bruijn* assemblers work best up to 50-80x coverage (probably even 100x [e.g. (Desai *et al*. 2013)] or 120x for heterozygous genomes (Vurture *et al*. 2017)), above which spurious contigs begin to appear due to the presence of more sequencing errors [e.g. (Desai *et al*. 2013; Mirebrahim *et al*. 2015; Lonardi *et al*. 2015)]. Therefore, trimming and coverage reduction have jointly contributed to obtain a much cleaner de *Bruijn* graph, and subsequently better super-reads to be aligned to the long reads. In fact, the contiguity of the super-reads built by MaSuRCA for pcla1 is lower than for pcla4. These super-reads have N50 538 bp and 573 bp, respectively.

The most contiguous hybrid assembly (pcla4) comprises 724.62 Mb and has contig N50 (ctgN50) 15.86 Kb and scaffold N50 (scfN50) 19.72 Kb. We found only one scaffold aligning to the mitochondrial reference with two large mitochondrial segments repeated. This scaffold was removed from the assembly and the alternative mitochondrial genome was kept separately. Our mitochondrial assembly matches the reference genome with 99.8% identity. In fact, the few differences are restricted to one indel and two Single Nucleotide Variants (SNVs) (Figure 5).

**Figure 4.**
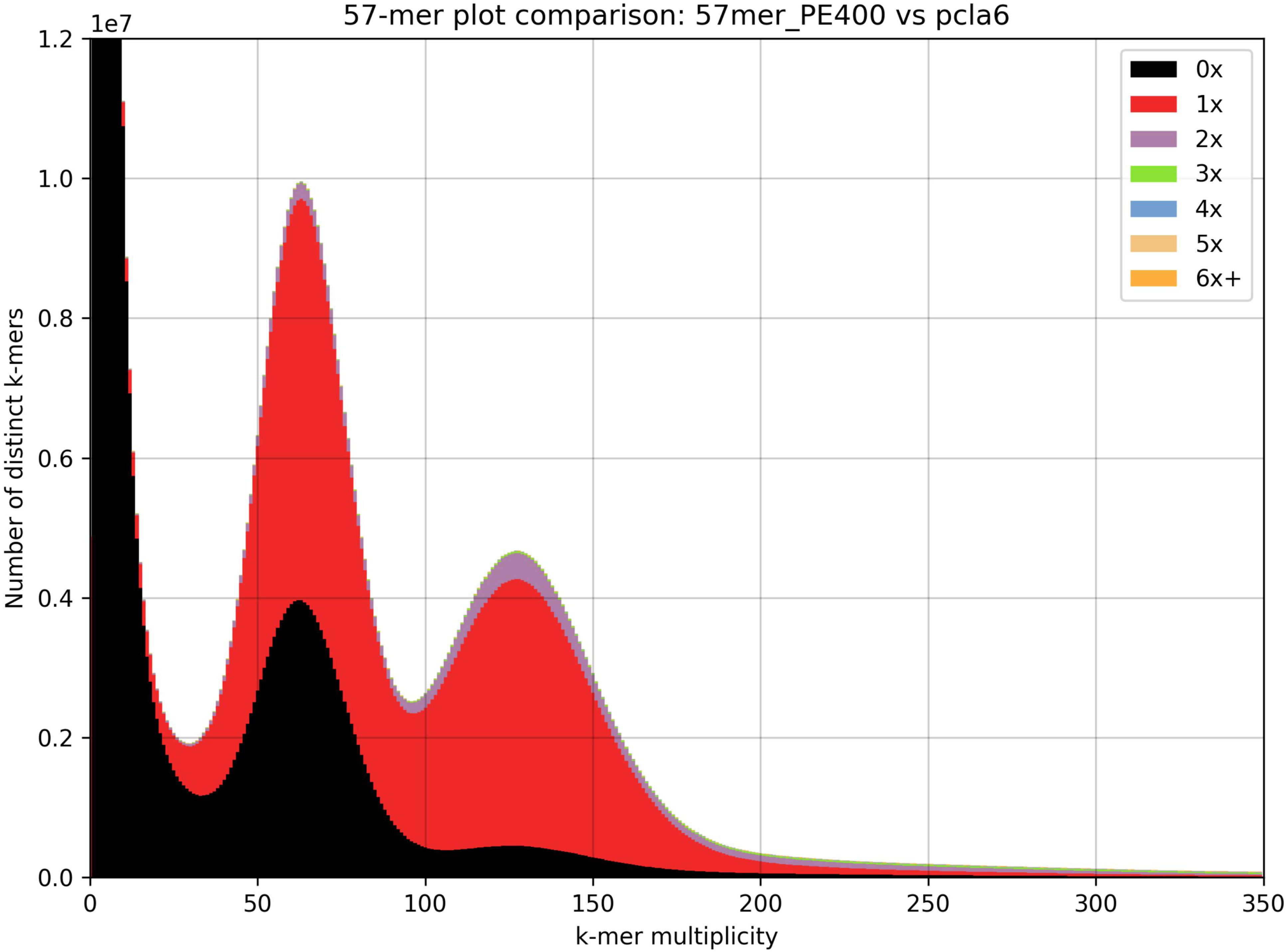
Comparison between the k-mer (k=57) spectra of PE400 and the *pcla*6 assembly. This is a stacked histogram produced with KAT that shows the spectra copy number classes along the assembly.

**Figure 5.**
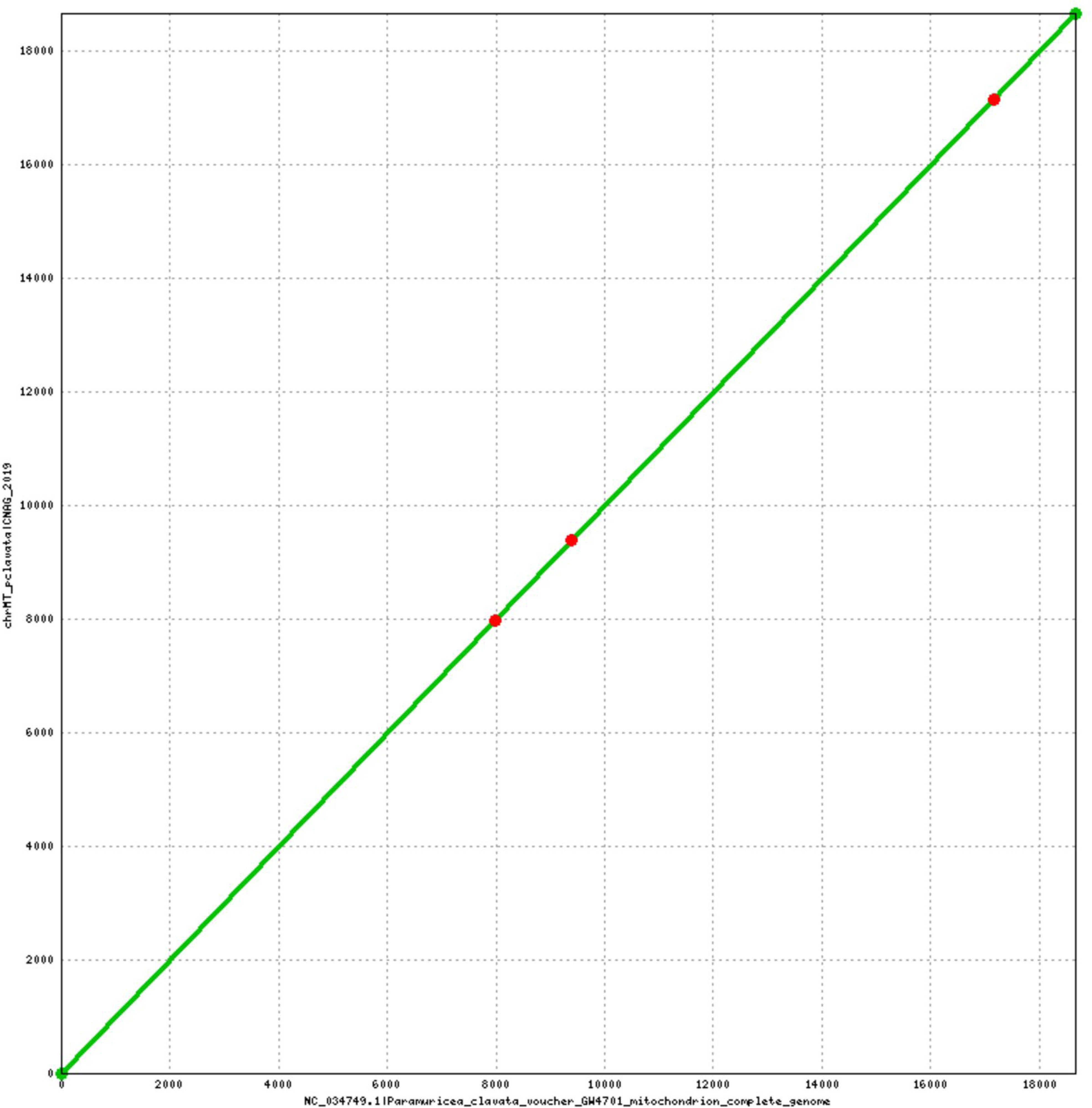
Alignment of the complete mitochondrial assembly against the NCBI reference genome (NC_034749.1) using DNAdiff v1.2 (MUMMER 3.22 package (Kurtz *et al*. 2004)). The figure produced with Mummerplot v3.5 (MUMMER 3.22 package) shows the location of the three mismatches found: one indel at position 9,389 plus two SNVs at positions 7,977 and 17,155, respectively.

**Figure 6.**
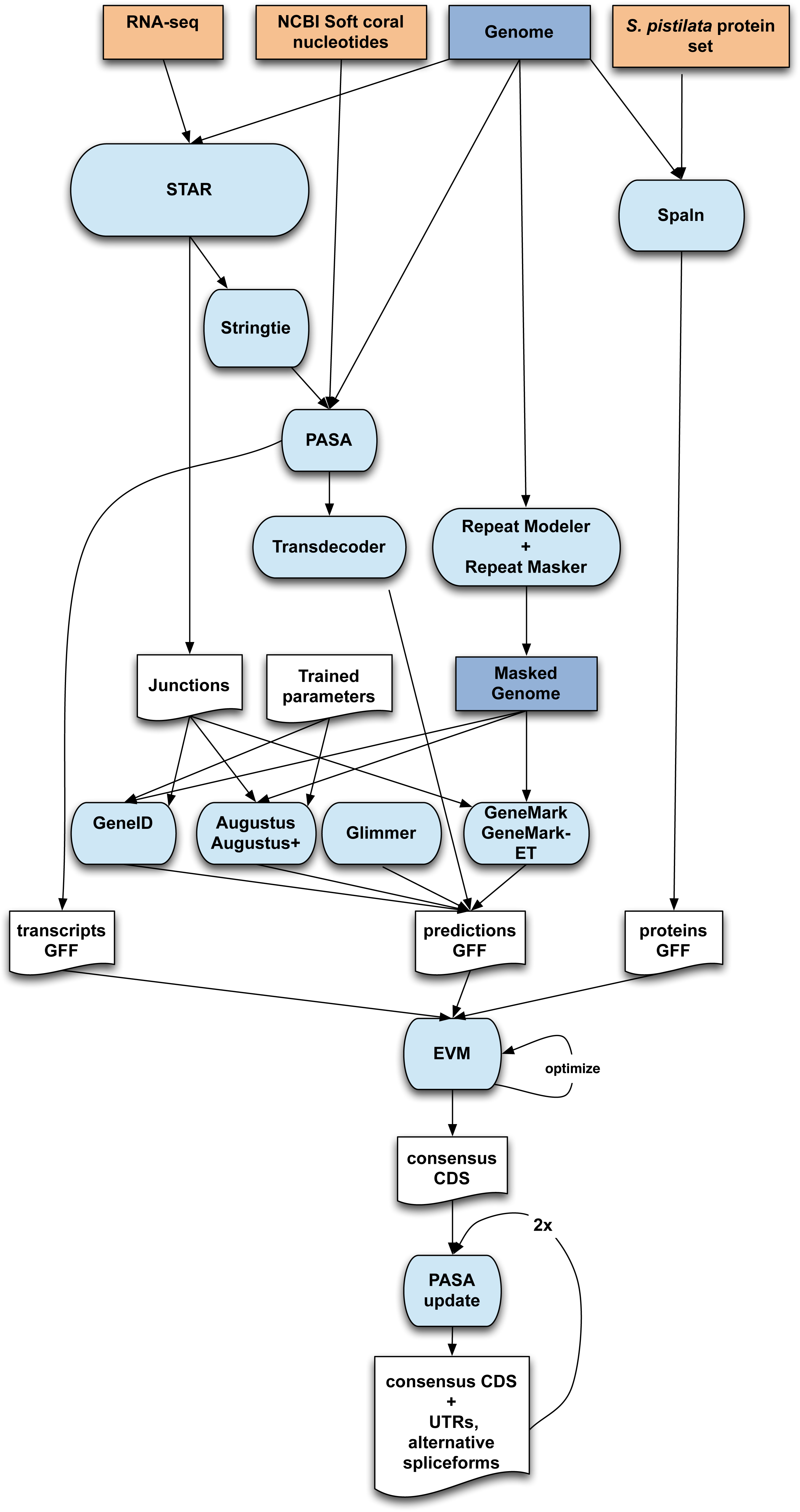
Genome annotation workflow.

Statistics of the decontaminated genome (pcla6) are shown in Table 4. Noticeably, the total contig length of pcla6, 711.53 Mb, is very close to the estimated genome size (711.53-712.31 Mb). However, a closer inspection of the spectra copy number (KAT (Marçais and Kingsford 2011), Figure 4), suggests that we failed to collapse some of the haplotypes (violet tip on top of the homozygous peak). On the other hand, owing to the use of long read data, it appears that true repeats have not been collapsed (bi-color tail evidencing the inclusion of 2x and 3x repeats). Consistent with the KAT results, the mapping rate of the Illumina PE400 against pcla6 is also very high, accounting for 98.33% of the total reads mapping to it.

**Table 3.** Sequence Contaminants.

| Species | % reads covered | No. reads covered | No. reads assigned | NCBI TaxID |
| --- | --- | --- | --- | --- |
| *Alteromonas mediterranea* * | 7.86 | 393,091 | 393,091 | 314275 |
| *Methanobacterium formicicum* | 0.47 | 23,547 | 23,547 | 2162 |
| *Candida dubliniensis* | 0.36 | 17,802 | 0 | 42374 |
| *Leishmania major* | 0.35 | 17,613 | 0 | 5664 |
| *Salmonella enterica* | 0.16 | 7,871 | 0 | 28901 |
| *Myceliophthora thermophila* | 0.15 | 7,378 | 0 | 78579 |
| *Streptococcus* sp. VT 162 | 0.14 | 6,921 | 6,921 | 1419814 |
| *Saccharomyces cerevisiae* | 0.1 | 5,153 | 0 | 4932 |
| *Theileria orientalis* | 0.07 | 3,293 | 0 | 68886 |
| *Thielavia terrestris* | 0.06 | 2,824 | 0 | 35720 |
| *Kluyveromyces lactis* | 0.04 | 2,018 | 2,018 | 28985 |
| *Human herpesvirus 7* | 0.03 | 1,475 | 1,475 | 10372 |
| *Cyprinid herpesvirus 1* | 0.03 | 1,282 | 1,282 | 317858 |
| *Mycobacterium kansasii* | 0.02 | 902 | 0 | 1768 |
| *Methanococcus voltae* | 0.02 | 787 | 0 | 2188 |
| *Tadarida brasiliensis circovirus 1* | 0.02 | 783 | 783 | 1732201 |

<sup>1</sup> The total number of read pairs screened was 5 million. <sup>2</sup> Actually included 10 matches to *A. mediterranea* (Taxonomy ID: 314275) and 393,081 matches to *A. mediterranea* U8 (Taxonomy ID: 1300257). Therefore, 393,081 matches represent 7.86% of the total reads.

**Table 4.** Contiguity of The Final Assembly (pcla6)

|  | Contigs |  | Scaffolds |  |
| --- | --- | --- | --- | --- |
|  | Length (bp) | Number | Length (bp) | Number |
| **N0** | 205,335 | 1 | 239,170 | 1 |
| **N5** | 76,693 | 356 | 90,053 | 305 |
| **N10** | 57,056 | 903 | 68,998 | 760 |
| **N15** | 46,501 | 1,596 | 56,084 | 1,336 |
| **N20** | 38,698 | 2,436 | 47,131 | 2,031 |
| **N25** | 32,714 | 3,440 | 40,326 | 2,847 |
| **N30** | 28,092 | 4,616 | 34,655 | 3,804 |
| **N35** | 24,351 | 5,979 | 30,123 | 4,907 |
| **N40** | 21,157 | 7,550 | 26,206 | 6,177 |
| **N45** | 18,348 | 9,358 | 22,716 | 7,636 |
| **N50** | 15,851 | 11,447 | 19,718 | 9,321 |
| **N55** | 13,636 | 13,868 | 17,012 | 11,266 |
| **N60** | 11,710 | 16,681 | 14,481 | 13,534 |
| **N65** | 9,939 | 19,981 | 12,273 | 16,205 |
| **N70** | 8,307 | 23,900 | 10,239 | 19,386 |
| **N75** | 6,790 | 28,637 | 8,329 | 23,246 |
| **N80** | 5,379 | 34,512 | 6,579 | 28,054 |
| **N85** | 4,053 | 42,119 | 4,926 | 34,299 |
| **N90** | 2,801 | 52,596 | 3,374 | 43,016 |
| **N95** | 1,476 | 69,693 | 1,776 | 57,221 |
| **N100** | 128 | 126,074 | 200 | 107,681 |
| **Total** | 711,495,393 | 126,074 | 712,409,171 | 107,681 |

In summary, the final assembly (pcla6) appears to be quite complete in terms of sequence and recovers a size that is very similar to the estimated by the k-mer analysis (Table 3, Figure 4). However, it is still highly fragmented with 107,681 scaffolds and scfN50 close to 20Kb. The reasons for this are likely the short length (N50=2.67Kb) and the sub-optimal coverage of nanopore reads: less than 10x (Zimin *et al*. 2017)).

### Genome annotation

The repeat annotation step results in 48.28% of the assembly identified as repeats, but this percentage rose to 49% after decontamination of the genome (see below). Table 5 shows the proportions of each repeat type for the pcla6 assembly (notice that they sum more than 49% as some positions fall in more than one category). The BUSCO analyses for gene completeness reports 77% complete genes and 8.9% fragmented genes. In total, the *Paramuricea clavata* genome assembly contains 76,508 annotated protein-coding genes, which produce 85,763 transcripts (1.12 transcripts per gene) and encode for 84,766 unique protein products. The annotated transcripts have 4.58 exons on average, with 62.9% of them being multi-exonic (Table 6). In addition, 58,498 non-coding transcripts have been annotated, of which 29,121 and 29,377 are long and short non-coding RNA genes, respectively. The high number of protein coding genes in comparison to other octocoral species, such as the sea pansy (Jiang *et al*. 2019) that has around 23,000 genes, is likely due to the high fragmentation of the genome. The facts that 38% of the annotated protein-coding genes contain only partial Open-Reading frames and that only 41% of the proteins have been functionally annotated support our previous statement.

**Table 5.** Repeat Annotation.

| **Repeat type** | **Bases covered** | **% genome** |
| --- | --- | --- |
| LINE | 38,376,659 | 5.39 |
| SINE | 11,263,955 | 1.58 |
| DNA | 75,455,215 | 10.60 |
| LTR | 10,131,293 | 1.42 |
| RC | 1,586,523 | 0.22 |
| Satellite | 1,785,201 | 0.25 |
| SnRNA | 189,245 | 0.03 |
| Simple repeat | 1,517,444 | 0.21 |
| Unkown | 226,267,576 | 31.80 |

**Table 6.** Genome Annotation Statistics.

|  | **Pacla6A annotation** |
| --- | --- |
| Number of protein-coding genes | 76,508 |
| Median gene length (bp) | 1,470 |
| Number of transcripts | 85,763 |
| Number of exons | 326,917 |
| Number of coding exons | 314,665 |
| Coding GC content | 41.65% |
| Median UTR length (bp) | 469 |
| Median intron length (bp) | 363 |
| Exons/transcript | 4.58 |
| Transcripts/gene | 1.12 |
| Multi-exonic transcripts | 62.9% |
| Gene density (kb) | 9.31 |

### Genome-wide Heterozygosity (SNVs) and Microsatellite Markers

The first estimate of the heterozygosity rate was 0.492% and defined as the total number of heterozygous SNVs (2,618,189) divided by the total number of callable sites (532,009,648 bp). Focusing on SNV variants at callable sites that do not contain artificially duplicated 57-mers, the heterozygosity rate was 0.513%. Thus, on average, we expect to find approximately 5 SNVs per Kb (or 5,130 per Mb) in this species.

Most of the microsatellite markers previously identified in *P. clavata* have been correctly assembled and located in our genome, as we were able to detect the repeat sequence between both primers (Table S1). However, we have not assembled the region that corresponds to Pcla-26 and the repeat unit of Pcla-20 seems to be (TA) instead of (TTAT). Also, Pcla-27 is found twice in the assembly. With the exception of Pcla-9, -14, -24, -25, 13 microsatellites were associated to protein sequences. While Pcla-a, -10, -12, -17, -20, -22, -28, -29 were entirely found within intronic sequences, Pcla-d, Pc3-81, Pcla-21, Pcla-23 and Pcla-27 show one or both primer(s) or even the repeat sequence within exonic sequences (see Table S1).

## CONCLUSION

The genomic and transcriptomic resources developed here will open up new ways to study on the ecology and evolution of *Paramuricea clavata* and related octocoral species. In particularly, we aim to characterize the genomic factors and eco-evolutionary processes involved in the differential responses to thermal stress observed during the warming-induced mass-mortality events. We are currently re-sequencing the genome of thermo-resistant and sensitive individuals identified through a transregional common garden experiment developed in the framework of the MERCES project (European Union’s Horizon 2020 research and innovation program http://www.merces-project.eu). The targeted individuals come from 12 populations from five distant regions in the Mediterranean, Adriatic and Eastern Atlantic. This work will be complemented by gene expression analyses at the transcriptomic level involving some of the re-sequenced individuals. In parallel, we are developing an holobionte approach and the genomic resources will be used in complement with microbiome analyses to reveal the temporal and spatial interactions between *Paramuricea clavata* and its associated micro-eukaryote and prokaryote symbionts. We also aim to refine our current knowledge regarding the evolutionary and demographic history of the species and are currently expanding our sampling to regions that were not covered by the common garden experiment (e.g. Turkey, Algeria, South West of Spain). In the end, the genome assembly described here will directly enable applications such as active restoration or assisted evolution (van Oppen *et al*. 2015) aimed at the conservation of *Paramuricea clavata*. Moreover, considering the structural role of *Paramuricea clavata* in biodiversity-rich coralligenous communities, future results obtained using the genomic and transcriptomic resources should extend to and benefit these communities as well.

## ABBREVIATIONS

MME: mass mortality events
Kb: kilobase pairs
bp: base pairs
BUSCO: Benchmarking Unifsal Single-Copy Orthologs
Gb: gigabase pairs
GC: guanine-cytosine
Mbp: megabase pairs
ONT: Oxford Nanopore Technologies
PE: paired-end.

## ACKNOWLEDGEMENTS

We thank the MERCES consortium, and particularly the members of the Work Package 3 (WP3-Restoration of coastal shallow hard bottoms and mesophotic habitats). We also would like to thank Drs. EA. Serrão and J. Boavida from the CCMAR (Centro de Ciencia do Mar, University of Algarve, Portugal) and Drs. S. Kipson and T. Bakran-Petricioli (University of Zagreb) for providing the samples used in the thermotolerance experiment. We are grateful to the staff of Experimental Aquarium Facilities at the Institute of Marine Sciences; E. Martinez and M. Delgado, for their technical support. We thank Oxford Nanopore Technologies for providing assistance with preparing and sequencing the libraries.

## FUNDING AGENCIES

We acknowledge the funding support of the European Union’s Horizon 2020 research and innovation program under grant agreement No 689518 (MERCES) and the Strategic Funding UID/Multi/04423/2013 through national funds provided by FCT – Foundation for Science and Technology and European Regional Development Fund (ERDF), in the framework of the programme PT2020.

JBL is funded by an assistant researcher contract framework of the RD Unit - UID/Multi/04423/2019 - Interdisciplinary Centre of Marine and Environmental Research – financed by the European Regional Development Fund (ERDF) through COMPETE2020 - Operational Program for Competitiveness and Internationalisation (POCI) and national funds through FCT/MCTES (PIDDAC). AA was partially supported by the FCT project PTDC/CTA-AMB/31774/2017 (POCI-01-0145-FEDER/031774/2017).

## AVAILABILITY OF DATA SUPPORTING

The assembly and raw reads have been deposited in the European Nucleotide Archive under the project accession PRJEB33489. The assembly, annotation, genome browser and BLAST server are also accessible via denovo.cnag.cat/pclavata.

## AUTHORS’ CONTRIBUTIONS

JBL and JG planned and granted the funding to start the project. JBL, JG, DGS, PLS and CL performed the sampling and the thermal stress experiment in aquaria. JBL, JG, FC, TA and MGu designed the sequencing strategy. JBL extracted the genomic DNA for the sequenced individuals. MGu performed the Illumina sequencing. RA and MGu performed the Nanopore sequencing. FC and TA performed the genome assembly. JGG and TA performed the genome annotation. FC performed the single-nucleotide variation analysis. JGG located the microsatellite loci in the final assembly. JBL, FC and JGG wrote a first manuscript draft. TA and MGu did a first revision of the original manuscript. All authors contributed to the writing of the supplementary data notes and to the preparation of supplementary tables and figures. All the authors read and approved the manuscript. MGu, TA and JG supervised the whole study.

---

**Table S1.**
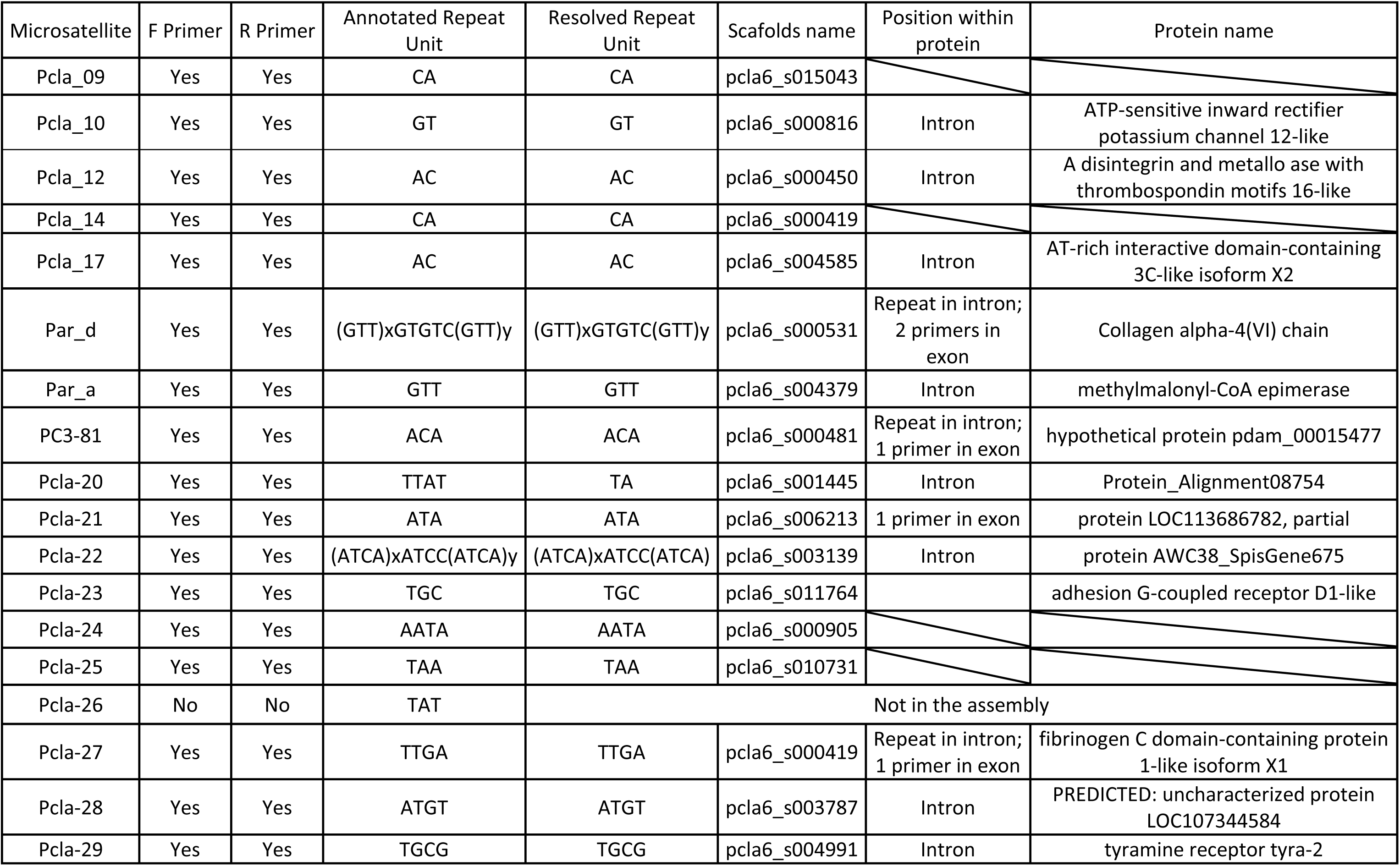
Mapping of microsatellites.

